# The fingerprint of functional strategies in Mediterranean seagrass fish assemblages

**DOI:** 10.1101/2025.03.13.642835

**Authors:** Bruno Bellisario, Azzurra Lattanzi, Roberta Cimmaruta

## Abstract

In the Mediterranean Sea, fish assemblages associated with seagrasses host a relevant percentage of the total fish biodiversity. While the structural complexity, coverage and distribution of seagrasses are known to influence species richness and abundance, their effect on the functional composition of faunal assemblages is still unclear. Moreover, little is known about the relationships of fish specific feeding, behavioural and reproductive strategies with the potential use of this habitat during different life stages. We characterized the functional structure of fish assemblages in three main Mediterranean seagrasses, using specific functional traits known to be associated with the potential use of meadows (e.g., as nursery, refuge and/or foraging areas). We also hypothesized that given combinations of traits could be associated with the probability of species being observed on seagrasses during well-defined life stages, thereby linking different strategies to habitat use. Fish functional assemblages are non-random structured, independently of either seagrass, taxonomic composition or species diversity, suggesting a convergence in their functional structures. Mediterranean seagrasses host a core of functionally similar fish species characterized by r-like reproductive strategies and low-to-intermediate trophic levels, although differing in the range of traits exhibited by predator fauna. Moreover, unique combinations of traits are significantly associated with the likelihood of fish being observed on seagrass during specific life stages. Our findings reinforce the view that the predominant use of seagrasses by fish may be driven by a trade-off between different life-history and feeding strategies. As functional strategies are largely involved in the structuring of food webs and species recruitment, their characterization in seagrass-associated fish assemblages would enhance our understanding of the impact of multiple threats on marine biodiversity.

## 1. Introduction

Seagrasses represent a key habitat in the marine environment and play a crucial role in maintaining marine biodiversity (Duffy, 2006). Indeed, seagrasses are habitat-forming species, facilitating the occurrence and persistence of other organisms by creating favourable conditions in terms of food supply and shelter from predation, ultimately influencing the distribution and abundances of species (Bertness & Callaway, 1994; Bulleri et al., 2018). This positive feedback is often associated with the ability of seagrasses to create complex structures and to form dense meadows with respect to bare substrates, although hosting less biodiversity than mangroves or coral reefs (Jones et al., 2021; Yeager & Hovel, 2017). Moreover, seagrasses play an important role in shaping fish coastal communities in terms of spatial connection among fundamental habitats such as mangroves, coral reefs and rocks, whose connectivity enhance fish diversity (Gilby et al., 2018; Goodridge Gaines et al., 2022; Unsworth et al., 2008).

Among the five native seagrasses of the Mediterranean Sea, *Posidonia oceanica* (L.) Delile, *Cymodocea nodosa* (Ucria) Ascherson and, to a lesser extent, *Zostera marina* are the most prevalent, differing in canopy complexity, distribution, and environmental tolerance (Ruiz et al., 2009). *P. oceanica*, the only endemic Mediterranean seagrass, forms dense, structurally complex meadows up to 40 m deep, spanning from the Alboran Sea to the Levantine Basin, although its distribution is declining due to anthropogenic impacts (Chefaoui et al., 2018; Ruiz et al., 2009). *C. nodosa* is widely distributed across the Mediterranean and adjacent Atlantic Ocean and tolerates broader environmental ranges and disturbances. This adaptability allows it to colonize degraded areas previously dominated by *P. oceanica* (Ruiz et al., 2009). In contrast, *Z. marina* has a restricted range, primarily in cooler northern regions such as the Adriatic Sea and western Mediterranean, where it forms sparse meadows in sheltered, shallow waters (Ruiz et al., 2009).

Besides differences in their distributional range, seagrasses in the Mediterranean Sea also differ in peculiar characteristics able to determine their structural complexity (e.g., canopy height, number of leaves per shoot, length and width of the leaves). For instance, *P. oceanica* forms dense, canopy-like meadows consisting of tall shoots (with leaves reaching up to 1.5 meters in length) and long-lasting matte due to its robust rhizome system. Conversely, *C. nodosa* creates looser, patchy beds, offering moderate habitat complexity, whilst *Z. marina* has only sparse shoots with thin leaves, providing basic habitat support in cooler or less stable zones (Ruiz et al., 2009). Such characteristics have been found to potentially determine the abundance and diversity of fish fauna (Lürig et al., 2016; McDevitt-Irwin et al., 2016). However, the extent to which the functional composition of fish assemblages differs between Mediterranean native seagrasses remains largely unexplored, with research primarily focusing on comparisons between single seagrass species *versus* other substrates, such as bare areas or macroalgae-covered habitats (Biagi et al., 1998; Bussotti & Guidetti, 2011; Franco et al., 2006; La Mesa et al., 2011).

A recent meta-analysis (Lattanzi et al., 2024) has shown how, in the Mediterranean Sea, seagrasses host about 38% of total fish species, with most of them (240 over 248) spotted on the endemic *P. oceanica* and 82 recorded as exclusive (i.e., never observed in other seagrasses). Interestingly, such exclusive species have been reported almost totally as adults, being in most cases large-sized, highly-mobile predators (Kalogirou et al., 2010; Lattanzi et al., 2024). Mean-sized and mobile predators as, for instance, the Atlantic horse mackerel (*Trachurus trachurus*), are indeed reported as occasional visitors on seagrass meadows, using this habitat exclusively for active prey foraging (Kalogirou et al., 2010). Another meta-analysis based on Mediterranean visual-census data has suggested that large predators such as the greater amberjack (*Seriola dumerili*) or the goldbloch grouper (*Epinephelus costae*) play a relevant role in shaping *P. oceanica* prey community, although being transient species not associated with the seagrass itself (Zubak et al., 2017).

How species perceive and use habitats may relate to peculiar adaptations linking species-specific attributes (e.g., morphology, ecological requirements, trophic ecology, behaviour) to the physical complexity of habitats, suggesting non-random species assembly along gradients of increasing/decreasing complexity (Ferreira et al., 2023; Sanabria-Fernández et al., 2024). Moreover, many marine organisms, in particular fish, often undergo changes in the diet, feeding modes, or social aggregation during the development, determining an ontogenetic shift in the use of habitats (Fobert et al., 2020; Polte et al., 2017). Therefore, it is possible that specific combinations of traits may influence the use of seagrass habitat by fish, with occasional visitors observed almost exclusively as adults having functional strategies different from resident species with a stronger tendency to use such habitats during their entire lifetime. How these specific functional strategies, determined by peculiar combination of both life-history and feeding traits, may be related to fish use of habitats during fish life cycle is a topic that has been recently object of attention by researchers (Koutsidi et al., 2020; Pecuchet et al., 2017). Indeed, community assembly mechanisms have been typically examined based on abundance data or taxonomic diversity, possibly hindering the relative importance of deterministic processes driving the convergence/divergence of traits among co-occurring species (Ford & Roberts, 2019; Kraft et al., 2015).

In this paper, we analysed the functional diversity of fish assemblages in three main native Mediterranean seagrasses, with the aim of identifying traits that may discriminate their communities and exploring the potential relationships between differences in the seagrass structural complexity and the functional traits of associated fish fauna. We also investigated whether the probability of species occurrence on seagrasses at different life stages is associated with unique combinations of traits that define specific life cycle and feeding strategies. To this end, we used an up-to-date and recently published database about the presence and the life stages at which fish species have been reported on Mediterranean seagrasses, alongside specific functional traits known to be associated with habitat use by fish. We discuss our results in light of the different mechanisms possibly associated with the structuring of fish communities, since either the importance of seagrass structural complexity in providing refuge and resources or the presence of predators in shaping fish communities have been alternatively proposed as main shaping forces (Heck & Orth, 1980; Schultz et al., 2009; Zubak et al., 2017).

## 2. Materials and methods

### 2.1 Data extraction and functional traits selection

We used a recently compiled database (Lattanzi et al., 2024) to retrieve all available information about the presence and the life stages at which fish species have been reported from the three main native seagrasses of the Mediterranean Sea, that is, meadows dominated by *P. oceanica, C. nodosa* and *Z. marina*. We checked for data consistency and excluded from the list those species for which we did not have at least two papers reporting the life stage. The most recent version of FishBase (Froese & Pauly, 2024), FishMed (Albouy et al., 2015) and a survey of current literature (last update 22/01/2025) were used to extract a set of specific functional traits related to the trophic ecology, size, growth and generation time (Table 1).

**Table 1.** List and definition of the functional traits used.

| Trait | Acronym | Type | Definition |
| --- | --- | --- | --- |
| Feeding behaviour | AMB | Factor | The main feeding behaviour of a species regardless of its trophic position. AMB - Ambush predators; AP - Active predators; GR - Grazers (feeding on substrates, either detritus, plants, animals and/or small crustaceans) |
|  | AP |  |  |
|  | GR |  |  |
| Maximum length | ML | Numeric | The size (in cm) of the longest individual recorded from a stock. A proxy for body size. |
| k | k | Numeric | The parameter of the von Bertalanffy growth function (also known as growth coefficient), expressing the rate (year<sup>-1</sup>) at which the asymptotic length is approached. Faster growth rates are associated with higher k values. |
| Generation time | Gt | Numeric | The time period from birth to average age of reproduction. |
| Trophic level | TL | Numeric | The position of a given species in the food chain. |

Indeed, specific combinations of life history and feeding-related traits might determine how species use habitats according to the peculiar characteristics of the seagrass (e.g., structural complexity, ability to form extensive meadows, distributional ranges), as either refuge, nursery and/or foraging areas. Before analysis, we controlled for multicollinearity using the eigenvalues of the predictor correlation matrix to measure the condition number (κ), which quantifies the sensitivity of the predictors (Belsley et al., 1980; Montgomery et al., 2021). Values of κ < 30 indicate a negligible or absent collinearity, while values of κ > 30 and κ > 100 indicate a moderate to severe collinearity between the selected functional traits (see Supplementary Information for further details).

### 2.2 Replication statement

| Scale of inference | Scale at which the factor of interest is applied | Number of replicates at the appropriate scale |
| --- | --- | --- |
| Ecological community | Mediterranean Sea | 169 (fish) + 3 (seagrasses) |

### 2.3 Functional traits imputation

To fill missing information in the functional traits of species, we performed a trait imputation procedure using random forest algorithms as in the function ‘impute’ from the *funspace* package of R (Carmona et al., 2024; Pavanetto & Puglielli, 2024; R Core Team, 2024). Previous works have shown how traits imputation can handle missing data effectively and can successfully quantify large-scale functional diversity even with high data missingness (Johnson et al., 2021; Stewart et al., 2023). Although its efficacy on categorical traits still remains controversial (Akande et al., 2017; Kim et al., 2018), the use of phylogenetic information (when available) may improve the performance of trait imputation, especially when traits have high levels of correlation and are conserved in the phylogenetic tree (Johnson et al., 2021; Pigot et al., 2020; Stewart et al., 2023). Moreover, the strong relationship between form and function in fish (e.g., body shape and size often relate with the feeding behaviour and/or life-history strategies) allows phylogenetically informed imputation methods to be particularly reliable for this group (Thorson et al., 2023).

Before analyses, we subdivided the dataset based on the main two classes of Actinopterygii (ray-finned fish, downloaded from https://fishtreeoflife.org) and Chondrichthyes (cartilaginous fish, Villalobos-Segura et al., 2022), for which complete phylogenetic information is available only separately. Then, we filled missing information by means of a phylogenetic trait imputation procedure on the two separate datasets, by adding the corresponding phylogenetic trees and finally merged the datasets. Quantitative traits were log-transformed, centred and scaled before imputation procedure. Following Penone et al. (2014) and Carmona et al. (2021), we added ten phylogenetic eigenvectors to the trait matrix, while species not included in the phylogeny were added to the root of their genus using the ‘addingSpecies’ argument of the above-mentioned ‘impute’ function.

Since a rigorous data checking for biases before and after imputation is necessary to minimize errors (Johnson et al., 2021), we controlled for the number of missing information in both each trait and combinations of traits by using the function ‘aggr’ in the package *VIM* of R (Kowarik & Templ, 2016). We further checked whether data were completely missing at random (MCAR), at random (MAR), and not at random (MNAR), as they might affect the reliability of imputation, especially in presence of MNAR-type data (Nakagawa & Freckleton, 2008). To this end, we used the Little’s test, a multivariate test based on maximum likelihood estimates to compare the observed variable means for each pattern of missing data with the expected population means (Little, 1988). A statistically significant result provides evidence against MCAR, suggesting that data might not be missing at random (MNAR). The Little’s test was computed using the function ‘na.test’ in the *misty* package of R (Yanagida, 2024). Post imputation diagnostic was evaluated using the function ‘post_imp_diag’ implemented in the package *missCompare* of R (Varga & Westergaard, 2024), a missing data imputation pipeline which gives an informative assessment of the congruence between original (with missing data) and imputed dataset via bootstrapping. See Supplementary Information for further details.

### 2.4 Functional structure

The multivariate space given by a set of traits determines the community functional space, and how densely species occupy different parts of this space defines the functional structure of a given community (Carmona et al., 2016). From a mathematical point of view, the functional structure can be measured with a traits’ probability density function (TPD) using a kernel density estimation with unconstrained bandwidth, starting from raw traits data or any given combination following a multivariate ordination (Carmona et al., 2019; 2021). According to Carmona and co-authors (2016), the functional space analysis provides useful information on both functional richness (FRic, the amount of functional space occupied by species) and divergence (FDiv, how much the species distribution is close to the extremes of the functional volume), as well as on the identification of areas of particularly dense species occupation in the traits’ space (i.e., functional hotspots, *sensu* Díaz et al., 2016 and Carmona et al., 2021). Given the mixed nature of traits used in this work (Table 1), we used a Factor Analysis of Mixed Data (function ‘FAMD’ in the package *FactoMineR* of R, Lê et al., 2008), an ordination technique that uses a principal components analysis (PCA) for quantitative variables and a multiple correspondence analysis (MCA) for qualitative variables (transformed into a disjunctive data table). The first two axes of ordination were used to measure the TPD (function ‘funspace’ in the package *funspace* of R) of fish assemblages in *P. oceanica, C. nodosa* and *Z. marina*.

We also tested whether the fish functional spaces measured in different seagrasses significantly deviated from randomness by comparing the amount of functional space occupied by the species against 100 null TPDs following a uniform distribution, where all combinations of traits within the range of observed functional trait space axes were equally possible (function ‘funspaceNull’ in the package *funspace* of R). The observed and null trait space were compared by means of a two-tailed *t*-test using standardized effect sizes (SES, the difference between the observed value and the mean of the simulated values after standardization by their standard deviations, Diaz et al., 2016).

### 2.5 Fish traits combination and life stages in *P. oceanica*

We explored the hypothesis that the presence of species on seagrasses in well-defined life stages might reflect the differential use of the habitat, likely due to species-specific combinations of traits. We hypothesized that those species commonly reported exclusively as juveniles will be more likely to use seagrass as nursery and/or refuge areas. Conversely, species reported exclusively as adults should represent occasional visitors, whose presence would be mainly linked to the use of seagrass as foraging area. Those species reported both as juveniles and adults would use the seagrass as either nursery, refuge and foraging areas, being classified as true seagrass residents. Unfortunately, information on the fish life stages is very scarce in the literature for *C. nodosa* and *Z. marina* (even if considering mixed meadows), so preventing to test this hypothesis for these seagrasses. We therefore focused this analysis on *P. oceanica*, for which a large amount of data is available.

To evaluate the relationships between the functional space of *P. oceanica* fish assemblage and life stage classifications, we followed Carmona et al. (2021, 2024) and: i) used generalized additive models (GAM) to model the relationship between the three life stage categories (adults, juveniles and adults + juveniles) with FAMD1 and FAMD2 and, ii) used the model fit to make predictions across the functional space grid and estimate how the response variable (i.e., life stages) varies across different functional trait combinations. Here, we used a slightly modified version of the original ‘funspaceGAM’ function in the *funspace* package of R, since it does not accept a multinomial family (a complete R code can be found in the Supplementary Information). We first fitted a Bayesian generalized additive model with smoothing splines using the *brms* package in R (Bürkner, 2017) and then used the fitted model to predict probabilities across the functional space. Specifically, we applied the ‘fitted’ function from the *brms* package to a new data grid representing the functional space derived from the traits’ probability density function (TPD, see the above section 2.4, Functional structure). This approach allowed us to estimate the probability of species being classified as either adults, juveniles or both (adults and juveniles) based on their functional trait positions. The results provided a probability surface reflecting how the likelihood of each life stage varied across the functional space. See Supplementary Information for further details.

## 3. Results

### 3.1 Fish functional structure comparison among seagrass habitats

From the total pool of 248 species already reported on seagrasses at Mediterranean Sea scale (Lattanzi et al., 2024), the data filtering for known life stages provided a total of 167, 123 and 36 species reported on *P. oceanica, C. nodosa* and *Z. marina*, respectively. Fish assemblages had a nested structure with respect to seagrasses, with species associated with *Z. marina* representing a subset of those observed in *C. nodosa* and these latter being a subgroup of the numerous species found in *P. oceanica*. For all seagrasses, the fish functional space captured more than 60% of total variation, with the first axis of ordination positively associated mainly with the maximum length and generation time (ML and Gt in Fig. 1 and Table 2). To enhance the clarity and accessibility of the results, we developed an interactive figure as a web application using Shiny (Chang et al., 2024). The application is accessible via the following link [https://brunobellisario.shinyapps.io/Fish4Fun/].

**Table 2.** Contribution of the fish functional traits to the first two axes (FAMD1 and FAMD2) of the Factor Analysis of Mixed Data (FAMD, see Materials and methods). Traits coded as in Table 1.

|  | *Posidonia oceanica* |  | *Cymodocea nodosa* |  | *Zostera marina* |  |
| --- | --- | --- | --- | --- | --- | --- |
|  | FAMD1 | FAMD2 | FAMD1 | FAMD2 | FAMD1 | FAMD2 |
| TL | 0.694 | 0.263 | 0.645 | 0.455 | 0.376 | 0.724 |
| ML | 0.841 | 0.169 | 0.868 | -0.215 | 0.901 | -0.003 |
| k | -0.648 | 0.567 | -0.676 | 0.199 | -0.686 | 0.494 |
| Gt | 0.839 | -0.330 | 0.865 | -0.050 | 0.887 | -0.264 |
| AMB | 0.474 | -1.027 | 0.477 | 3.150 | -1.382 | -2.636 |
| AP | 1.391 | 1.454 | 2.039 | -0.346 | 2.050 | 0.944 |
| GR | -0.608 | -0.339 | -0.646 | -0.299 | -0.463 | -0.142 |
| % of variance | 43.205 | 19.064 | 46.498 | 19.093 | 43.477 | 19.430 |

**Figure 1.**
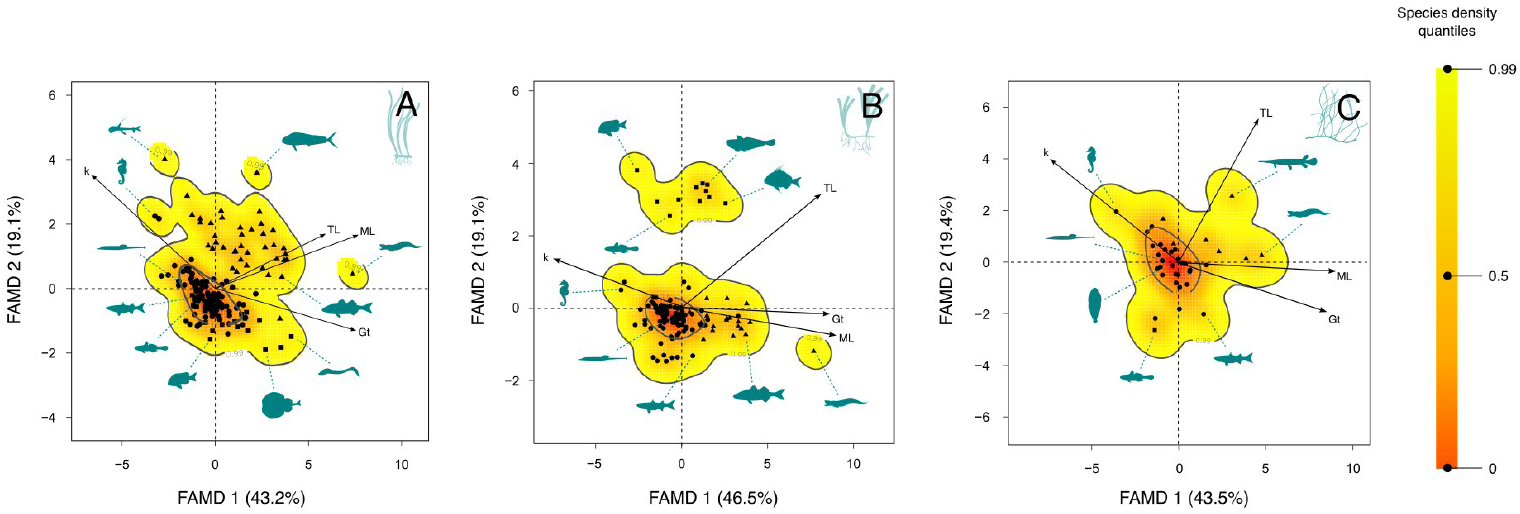
Functional space of fish assemblages in *Posidonia oceanica* (A), *Cymodocea nodosa* (B) and *Zostera marina* (C), showing the probabilistic species distributions in the spaces defined by the first two dimension (FAMD 1 and FAMD 2) of the Factorial Analysis of Mixed Data (FAMD, see Materials and methods). Arrows indicate the direction and weighting of quantitative traits (TL = trophic level; ML = maximum length; Gt = generation length and k = growth coefficient) in FAMD (see Table 1). Symbols indicate the feeding behaviour of species (qualitative trait): circles = grazers; triangles = active predators; squares = ambush predators (see main text). The colour gradient indicates the density of species within the functional space (increasing density from light yellow to red), while thick contour lines indicate the 0.5 (i.e., functional hotspots, see main text) and 0.99 quantiles of distributions. Fish silhouettes were downloaded from PhyloPic (https://www.phylopic.org) under Creative Commons licenses. An interactive version of this figure can be viewed at [https://brunobellisario.shinyapps.io/Fish4Fun/].

In *P. oceanica*, the trophic level (TL) was strongly correlated with ML (*R*^2^ = 0.485, *p* < 0.001), being also an important trait determining the ordination of species along the first dimension of the FAMD (Fig. 1A and Table 2). Conversely, TL was only weakly, although significantly, correlated with ML and Gt in *C. nodosa* (*R*^2^ < 0.3, *p* < 0.05 in both cases), while showing no significant correlation in *Z. marina* (*R*^2^ < 0.2, *p* > 0.09 in both cases).

However, the trophic level TL contributed equally to the distribution of species along both axes of ordination in *C. nodosa* and almost equally in *Z. marina* (Fig. 1B and 1C and Table 2). Both the growth coefficient k and feeding behaviour contributed to the ordination of species along the first two axes, with the only exception of *C. nodosa*, where the feeding strategy was the main trait determining the distribution of species along the second axis (Fig. 1B and Table 2). Independently of the seagrass considered, the distribution of species within the measured functional spaces was uneven, with large species at the top of the food web and with feeding habits based on waiting and/or chasing prey always placed at the periphery of the functional space (Fig. 1). However, there were substantial differences among the seagrasses. Indeed, while in *P. oceanica* and *Z. marina* we observed a continuum in the functional space (with the only exception of three active predators in *P. oceanica*, Fig. 1A), in *C. nodosa* small-sized ambush predators represented a well-defined and separated cluster of species (Fig. 1B).

In all seagrasses, the functional hotspot (i.e., the smallest portion of functional space including 50% of the species, Diaz et al. 2016) consistently had a small extent if compared to the total area of the functional space, ranging from ca. 9% in both *P. oceanica* and *C. nodosa* to ca. 12% in *Z. marina* and included almost exclusively omnivore species, grazing on substrates mainly represented by seagrass leaves (Fig. 1). Overall, the position of large predators, far from the hotspots and peripheral in the functional spaces, indicated a low probability of occurrence of their trait’s combinations, while their scattered distribution suggested low or no functional redundancy. The peripheral position of the functional hotspot in *P. oceanica* and *C. nodosa* (closer to the boundaries of occupied space) indicated high values of functional divergence in these assemblages (FDiv = 0.65 and FDiv = 0.71, respectively), while in *Z. marina* the central position of the hotspot within the functional space suggested low functional divergence (FDiv = 0.482). The functional richness (i.e., the total amount of functional space occupied by the species) also differed between seagrasses and was higher in *Z. marina* and *P. oceanica* (FRich = 47.772 and 41.137, respectively) than in *C. nodosa* (FRich = 32.901).

Under the null scenario of a uniform distribution of traits among species (i.e., all trait combinations were equally possible for all species), the amount of functional space occupied was significantly smaller than the null expectation in all seagrasses, in line with the lumped distribution of species and the high functional divergence measured. Such pattern was more pronounced in *P. oceanica* and *C. nodosa* (*SES* = - 20.47 and *p* < 0.01 in both cases), than in *Z. marina* (*SES* = - 3.72 and *p* < 0.01).

### 3.2 Functional strategies and association with seagrasses

The multinomial regression model revealed significant associations between predictors (FAMD1 and FAMD2) and the likelihood of species being classified in the corresponding life stage categories (Table S1). All parameters showed strong convergence diagnostics with high effective sample sizes (Table S1), indicating robust and reliable posterior estimates (see Supplementary Information for more details). The combination of traits considered predicted an overall low probability of species to be present in *P. oceanica* exclusively as juveniles (Fig. 2 and Fig. S4).

**Figure 2.**
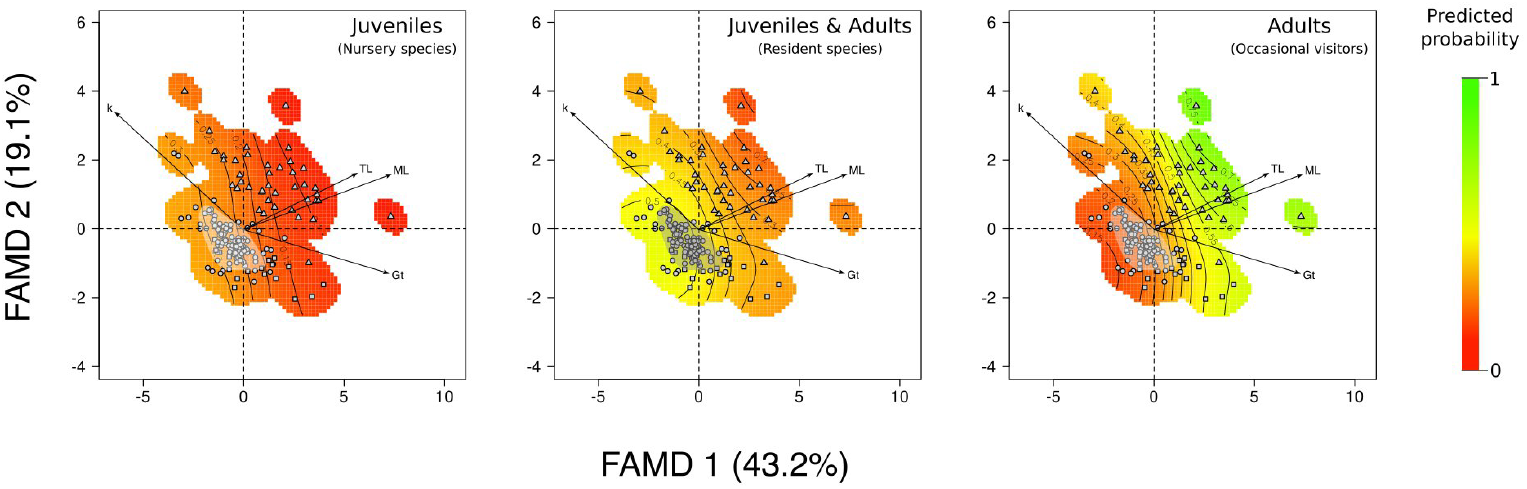
Probability distribution surfaces showing the likelihood of each life stage across the functional space of fish assemblage in *Posidonia oceanica*. The figure shows the probability of species being classified as either adults, juveniles and both (adults and juveniles) based on their functional trait positions according to a multinomial Bayesian generalized additive model (see section 2.5 in Materials and methods and Supplementary Information). Contour lines indicate the probability values of the distribution while the shaded area indicate the functional hotspot (see section 2.4 in Materials and methods).

Small-sized species with fast reproductive strategies and non-active predation modes had the highest probability to be observed as juveniles only, although this probability is very small (∼ 0.3, Fig. 2 and Fig. S4). Conversely, large-sized species at the top of the food web and with active predation modes had a higher probability (> 0.65) to be observed exclusively as adults (Fig. 2 and Fig. S4). Examples are large active and/or ambush predators as the common dolphinfish (*Coryphaena hippurus*) or the marbled electric ray (*Torpedo marmorata*) that have been reported in the literature as occasional visitors on *P. oceanica* (Kalogirou et al., 2010).

Species with intermediate combinations of traits as, for instance, medium size, low-to-intermediate trophic level and/or relatively high k had the higher probability to be present as both adults and juveniles (Fig. 2 and Fig. S4). Examples are the salema porgy (*Sarpa salpa*), the main consumer of *P. oceanica* in the Mediterranean Sea, or the spinefoots (*Siganus luridus* and *S. rivulatus*) that, despite being Lessepsian migrants, over the decades have established stable populations of residents on *P. oceanica* meadows in the eastern Mediterranean (Kalogirou et al., 2012). Furthermore, our findings showed that all species within the functional hotspot were characterized by having a relatively high probability (> 0.5) of being found as both adults and juveniles (i.e., resident species) if compared to other categories (Fig. 2 and Fig. S4), suggesting that the use of seagrass habitats may extend beyond being solely nursery or foraging areas, with a more complex interplay of functions likely coexisting.

## 4. Discussion

The taxonomic richness of fish assemblages differs in Mediterranean autochthonous seagrasses, as shown by a recent review (Lattanzi et al., 2024). In this work, we show how the three main native Mediterranean seagrasses host an array of functionally similar fish species, regardless of species richness. Fish assemblages in different seagrasses were characterized by non-random traits assembly, with small-to-medium fish mainly grazing on substrates representing the core of functionality in seagrass habitats. Conversely, large active (top) predators characterized by slow growth rates and high generation time were always found far from the core and at the periphery of the functional space, suggesting that such functional strategies are quite uncommon. Our results also indicate that peculiar combinations of both life-history and feeding strategies are related to the life stage at which species use the seagrasses, suggesting that functional traits may shape the ontogenetic habitat use of seagrass by fish.

### 4.1 Functional space comparison

Seagrasses are known to host a remarkable number of fish species in all the seas where they live and many studies have shown the importance of their structural complexity and coverage in structuring fish assemblages, evidencing that different seagrasses may provide different habitat features for associated species (Jones et al., 2021; Unsworth et al., 2022). Higher canopy height, greater number of leaves per shoot and shoot density increase the structural complexity of some seagrasses, allowing for a greater range of micro-habitats and faunal assemblages with greater abundance and richness (Jones et al., 2021). For instance, along the south-eastern coasts of Australia *Posidonia sinuosa* meadows form uniform and dense canopies inhabited by more fish species, with greater density and biomass with respect to the less structured meadows of either *Amphibolis griffithii* or *P. coriacea* (Hyndes et al., 2018).

In the Mediterranean Sea, the same pattern is observed, with native seagrasses hosting a significant percentage of the total fish diversity and sharing a large number of species. However, the fish species richness differs in meadows dominated by different seagrasses, with *P. oceanica* meadows hosting a unique and diversified fish fauna if compared to *C. nodosa* and, even more markedly, to *Z. marina* (Lattanzi et al., 2024). This is in line with seagrasses habitat complexity and range extension. The eelgrass *Z. marina* is a species well-adapted to colder temperatures and its distribution becomes sparse in the Mediterranean Sea, where it is mostly found as small, isolated stands and only occasionally occurring as dense beds (Ruíz et al., 2009). Conversely, *C. nodosa* is a warm tolerant species widely distributed throughout the Mediterranean Sea, where it forms large meadows sometimes in association with *P. oceanica*. This latter is the only endemic seagrass of the Mediterranean Sea where it forms dense and structurally complex meadows (Ruíz et al., 2009). These structural differences are due to both the larger size of *P. oceanica* and the more marked seasonal cycle of both *Z. marina* and *C. nodosa*, whose shot density strongly decreases during the cold season making the canopy poor or even absent (Guidetti & Bussotti, 2000; Marbà et al., 1996).

Our findings evidence that, despite the difference in species richness and meadow’s structure, fish assemblages in the three main Mediterranean seagrasses show a common core of functional strategies. Fast growth rates, small sizes and low trophic levels are the trademarks of fish functional traits in all studied seagrasses, shared by those species included in or closed to the functional hotspots, whose small space extent witnesses a high redundancy for these combinations of traits (Carmona et al., 2019; 2021).

In *P. oceanica*, functional core species are predominantly grazers with diverse diets resulting in low to moderate trophic levels, mainly consuming detritus, algae, and small crustaceans. This aligns with studies showing that Mediterranean seagrass-associated fish exhibit varied feeding modes and diets and are able of exploiting both benthic and planktonic resources, although being often classified as carnivores (Stergiou & Karpouzi, 2002; Vizzini, 2009). For instance, benthic feeders like *Diplodus sargus* and *D. annularis* also consume plankton, while typically planktivorous species like *Atherina presbyter* also feed on benthic invertebrates (Pinnegar & Polunin, 2000). Many core species overlap across the studied seagrasses, highlighting a consistent pattern, and include some of the most abundant species usually reported in the literature (e.g., sparids as *Boops boops* and seabreams, *Syngnathus* pipefishes, *Sparisoma cretense* and *Mullus surmuletus*, Guidetti & Bussotti, 2000; Máñez-Crespo et al., 2022). Ambush predators like Scorpaenidae and Gobiidae (e.g., *Scorpaena porcus, Gobius cruentatus, G. buchicchi/incognitus and G. cobitis*) belong to fish families frequently signalled in seagrass worldwide (Espino et al., 2015; Pollard, 1984). These species are recorded at the periphery of the functional space in all the seagrasses, particularly in *C. nodosa* and *Z. marina*. In *Z. marina*, ambushers account for a small percentage of species records, while in *C. nodosa* ambush predators are grouped into a distinct and well-separated cluster away from the functional hotspot. This suggests that, regardless of size and life history strategies, foraging modalities based on waiting and ambush represent uncommon strategies in both these seagrasses. Accordingly, the most abundant species recorded on *C. nodosa* in both Mediterranean and Atlantic areas by a recent study do not include ambushers, despite the meadows in Canary Islands benefit from local climatic stability and high temperatures. This finding suggests that the probability of observing this combination of traits is more likely determined by seagrass characteristics than by environmental drivers (Máñez-Crespo et al., 2022). Life history and feeding traits associated with larger body sizes, higher trophic levels, and active predation modes occupy the periphery of functional spaces, with a scattered distribution and far from the hotspot. This pattern indicates an overall low level of redundancy of this traits combination and evidence that species bearing these traits have lower probabilities of being observed on seagrasses. Examples are pelagic predators like *Euthynnus alletteratus, Auxis rochei, Trachurus mediterraneus, Sphyraena sphyraena*, and *S. viridensis*, often reported as occasional visitors on seagrass meadows (Guidetti & Bussotti, 2000; Kalogirou et al., 2010).

Functional richness and divergence are unrelated to taxonomic richness and vary across fish assemblages in the three seagrasses. The lumped distribution of species within the functional hotspot in *P. oceanica* and the discontinuity observed in the functional space of *C. nodosa*, both indicate a higher functional divergence compared to fish assemblages in *Z. marina*. Interestingly, this latter shows the higher functional diversity although hosting about a quarter of the species reported in other seagrasses, suggesting a high degree of complementarity in the functional strategies of associated fish fauna. Fish assemblages from *P. oceanica* are characterized by high values of both functional richness and divergence. Three species create a discontinuity in the functional space, without however influencing the global functional richness and divergence: the European conger (*Conger conger*), the garrick (*Cyclothone braueri*) and the common dolphinfish (*Coryphaena hippurus*). These species are all characterized by mesopelagic/pelagic distributional ranges or different habitat preferences, so that they are only occasionally reported on seagrasses (Álvarez et al., 2015).

Thanks to their structural complexity, seagrass habitats have often been proposed as a refuge for mobile demersal species, offering protection against predation, by reducing prey visibility and impeding predator movement, and reducing the risk of predation respect to surrounding open habitats (i.e., the so called ‘seagrass superiority hypothesis’, Heck & Orth, 1980). However, if this hypothesis successfully explains the high diversity and abundance of generalist species placed at intermediate to low trophic levels, it fails to account for the presence of large active predators that can increase mortality risks. The so-called ‘predation mode hypothesis’ considers this finding and suggests the predominant role of piscivorous predators in structuring fish communities in seagrass meadows (Kruschel & Schultz, 2020; Schultz et al., 2009; Zubak et al., 2017). Indeed, active predators may face challenges when foraging in dense seagrass meadows that can obstruct their movement and reduce visibility, while ambushers may prefer vegetated bottoms offering an optimal cover (Kruschel & Schultz, 2020; Schultz & Kruschel, 2010). Studies on Mediterranean fish assemblages are still poor, but experimental data from the Adriatic Sea have evidenced that aggressive small piscivorous and their fish preys tend to separate in shallow coastal habitats including *P. oceanica* and *C. nodosa* meadows, besides rocks and macroalgae, with preys preferring more homogeneous habitats and aggressive predators preferentially patrolling habitat edges (Kruschel & Schultz, 2020).

Our results are consistent with the proposition that seagrasses may virtually serve at the same time as a critical refuge for low-sized generalist species and as foraging habitat for a lower number of large piscivore predators that are occasional visitors. Active predation requiring stalking and attack is an established strategy in all seagrasses, while ambush predation is only poorly represented in the fish assemblage of *Z. marina*. According to the predation mode hypothesis this finding could be related to the different complexity of these habitats, with active predators less affected by the complexity of seagrass canopies due to their ability to patrol from neighbouring habitats and to attack specimens moving outside the meadow. Conversely, ambush predators are more closely related to the structural characteristics of seagrasses because they require adequate hiding places to attack their preys, which are more frequent in larger seagrass forming more complex habitats as *P. oceanica* and *C. nodosa* and almost lacking in poorly complex meadows of *Z. marina* (Horinouchi et al., 2009; Laurel & Brown, 2006; Schultz et al., 2009).

### 4.2 Functional strategies and life stages in *P. oceanica*

The use of seagrass habitats by fish may vary depending on species’ ontogeny, with some species present exclusively during a specific life stage (i.e., larvae, juveniles and/or adults) (Kalogirou et al., 2010). The distribution of highly mobile animals such as fish reflects the balance among conflicting demands associated with foraging and predation avoidance, often bringing animals to shift habitats because foraging needs, predation risks and optimal reproductive conditions may change during ontogeny as, for instance, due to increase in body size (Dahlgren & Eggleston, 2000; Fobert et al., 2020).

Our findings suggest that peculiar combinations of traits are significantly related to the probability of observing specific life stages in fish inhabiting *P. oceanica* meadows. Functional traits associated with fast reproductive strategies (e.g., small sizes, high growth coefficient, low generation time and trophic level) and grazer-like feeding behaviours are observed in species that have the highest probability of being found in *P. oceanica* as both juveniles and adults, therefore suggesting the use of this habitat during their entire life cycle both as refuge/nursery and foraging area. These traits match with those characterizing the species within the functional hotspot, indicating that the most common observed combinations of traits are also those allowing fish individuals to use the seagrass during all life stages, possibly increasing the shelter/nursery value of the habitat. Accordingly, most of the species within the functional hotspot are known to have an all-life-cycle use of the seagrass and are recognised as resident species strongly associated to this habitat: *Symphodus mediterraneus, S. ocellatus, Serranus cabrilla, Diplodus annularis, Spondiliosoma cantharus* and *Sarpa salpa* among others (Guidetti & Bussotti, 2000; Kalogirou et al., 2010).

On the other hand, large predators with slow reproductive strategies (i.e., high generation time and low growth coefficient) have nearly no probabilities to be observed in seagrass habitats during their juveniles’ stages, especially if characterized by an active predation mode, for which they may therefore represent mainly foraging grounds. This finding confirms that large pelagic species recorded in this habitat are transient species, spending their juvenile stage in other habitats and occasionally transiting on *P. oceanica* meadows (Jones et al., 2021). The same is observed for large ambushers, suggesting that size plays a prominent role in determining which species may utilize the meadows during the adult stage. This finding agrees with a large study carried out in eastern Mediterranean, showing that half of the 88 species sampled on *P. oceanica* along a whole year were occasional visitors, including some of the species here evidenced as large predators recorded only in their adult stage (e.g., the above mentioned grater amberjack *S. dumerili*, the Mediterranean morray *M. helena* or the marbled electric ray *T. marmorata*) (Kalogirou et al., 2010). These results also indicate that in *P. oceanica*, and possibly in other Mediterranean seagrass systems, large predators do not operate an ontogenetic shift using seagrass as nursery and leaving the meadows once reached a given size, as recorded for example in tropical seagrasses (Dorenbosh et al., 2005; Nakamura et al., 2012). Although limited to *P. oceanica*, the observed similarities in the functional spaces possibly suggest the same pattern for other seagrasses, because of the observed convergence in the functional strategies determining the seagrass use by fish.

## 5. Conclusions

By studying the functional space of seagrass-fish assemblages we identified a common ‘fingerprint’ in the functional strategies of associated fish species, highlighting that these assemblages are primarily shaped by a trade-off between various life history and feeding strategies. These strategies influence how species use habitats at different life stages, which in turn affects food web structure and species recruitment—both of which are essential for maintaining marine biodiversity. Functional diversity of seagrass-fish assemblages is an important aspect in marine biodiversity conservation since studies have shown how r-K selection strategies might have a role in determining the responses of species to disturbances, with r-selected species strongly affected by climate change but less impacted by fishing pressure, while the opposite is true for K-selected fish species (Graham et al., 2011; Lynam et al., 2017). Moreover, recent studies demonstrated how seagrass meadows in the Mediterranean Sea will experience a transition from long-lived, large and slow-growing species, such as *P. oceanica*, to small and fast-growing seagrasses as, for instance, the invasive *Halophila stipulacea*, as climate change progresses (Beca-Carretero et al., 2024). Overall, the identification of a ‘functional fingerprint’ in seagrass-associated fish assemblages would provide a useful baseline to understand how such transition will affect fish assemblages, improving our understanding on the effects of multiple threats on marine biodiversity.

