## Supplementary Information for "The fingerprint of functional strategies in Mediterranean seagrass fish assemblages"

Bruno Bellisario[1], Azzurra Lattanzi, Roberta Cimmaruta
Department of Ecological and Biological Sciences, Tuscia University of Viterbo,
Largo dell'Università Snc, 01100, Viterbo (Italy)

### Functional traits imputation

#### *Data inspection*

Before analysis, we controlled for multicollinearity using the eigenvalues of the predictor correlation matrix to measure the condition number ($\kappa$), quantifying the sensitivity of the predictors. Values of $\kappa < 30$ indicate a negligible or absent collinearity, while values of $\kappa > 30$ and $\kappa > 100$ indicate a moderate to severe collinearity. Here we found a value of $\kappa = 8.505$, meaning that the selected functional traits are well-conditioned (i.e., minimal or no multicollinearity), and the linear dependencies among them are not likely to distort subsequent functional analyses.

The number of observations (i.e., species) with complete functional information was 77%, with a total fraction of missingness of about 5%. Maximum length had complete observations for all species and, overall, quantitative variables had low percentages of missing data (trophic level, 0.59%; k and generation time, 1,18%). As expected, the only qualitative variable, that is, the feeding behaviour, had the highest percentage of missing data (ca. 22.5%). Overall, the Little's test was not significant ($n = 169$, $\chi^2 = 20.89$, $df = 13$ and $p = 0.075$), suggesting randomness in data deficiency and, therefore, excluding biases in no-data distribution between species and functional traits.

#### *Post imputation diagnosis*

Post imputation diagnosis showed an overall agreement between original and imputed values of traits (with the exception of maximum length for which we had no missing data), as observed by both the distribution of original and imputed values (Fig. S1) and the Kolmogorov-Smirnov test statistic D values, approaching zero for each imputation.

[1] Corresponding author

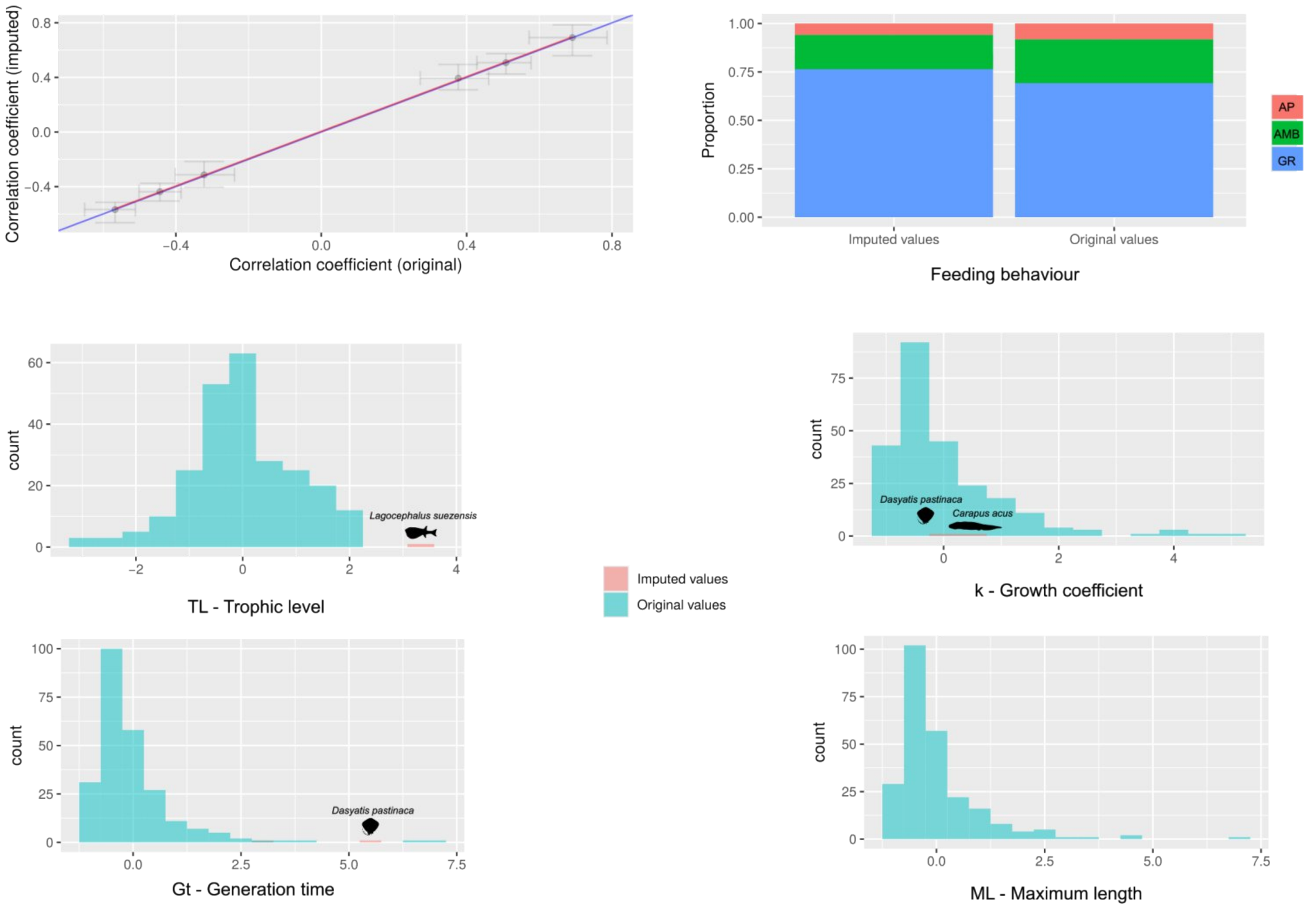


**Figure S1** – The correlation plot (top-left panel) shows the bootstrapped correlation coefficients from original and imputed data after 1,000 iterations. For a good imputation procedure, bootstrapped correlation coefficients (dots with ± standard error and red line) should align as much as possible to the blue line representing a perfect correlation between data, with intercept 0 and slope 1. The top-right panel shows the goodness of imputation procedure for the qualitative variable feeding behaviour, expressed as the proportion of samples (species) falling within each category before and after imputation: AP, active predators; AMB, ambush predators and GR, grazers (see main text for more details). Other figures show the overlaid distributions of original and imputed values for quantitative variables. Silhouettes show the species (or group of species) for which the functional traits have been imputed. Note that data are scaled to a 0-1 mean and standard deviation, respectively.

**Bayesian multinomial model with smoothing splines**

We modelled the categorical response variable, i.e., fish life stages (three categories, adults, juveniles and adults + juveniles, hereafter defined as 'both' or 'resident'), as a function of the two latent dimensions from the Factor Analysis of Mixed Data (FAMD1 and FAMD2, see main text) using a Bayesian generalized additive model with smoothing splines. The model was implemented using the *brms* package in R (Bürkner 2017), using 'both' as the baseline category (i.e., reference category). Smoothing splines were applied to both dimensions of the FAMD to model potential non-linear effects, while the Bayesian framework enabled estimation of posterior distributions for all parameters. Bayesian generalized additive models provide several advantages over traditional frequentist approaches, allowing for robust estimation of uncertainty through posterior distributions and credible intervals that are more interpretable than frequentist confidence intervals. Additionally, the framework enables the inclusion of prior information, which can help regularize models or incorporate existing knowledge about parameters and accommodates hierarchical structures and complex random effects, which are essential for modelling grouped or nested data. The response variable was modelled using a categorical family with a multinomial logit link, and separate smooth terms were fitted for each response category.

Model inference was based on 4 Markov Chain Monte Carlo (MCMC) chains, each with 2,000 iterations, including 1,000 warm-up iterations, resulting in 4,000 total post-warm-up draws. Model convergence was assessed using the potential scale reduction factor (aka Gelman-Rubin diagnostic) $\widehat{R}$, which indicates how well the chains have mixed, with values close to 1 suggesting proper convergence. Additionally, effective sample sizes (ESS) were examined to ensure sufficient independent samples. Model fit and performance were evaluated through posterior predictive checks and visual inspection of trace plots to confirm adequate sampling behaviour and distributional assumptions.

*Priors setting*

The intercept priors were specified to reflect the assumption that, before observing the data, the probability of a given life stage (adult or juvenile) is equally likely relative to the reference category (i.e., species found as both adults and juveniles aka resident species). These priors allow for moderate variability around this central expectation, meaning that while small deviations are likely, extreme deviations are less probable. This setup ensures that the model does not favour one life stage over another in the absence of strong evidence from the data, maintaining a balanced and flexible starting point for estimation. To incorporate biological expectations into the model, we assigned informative priors to the regression coefficients of the functional axes (FAMD1 and FAMD2) for each life stage. For adults, we specified priors centred at 1 with a standard deviation of 0.5, assuming a positive association between these functional space dimensions and the probability of being classified as an adult relative to resident species. For juveniles, we set priors centred at -1 with the same standard deviation, reflecting an expected negative association between these axes and juvenile classification compared to residents. These priors introduce moderate constraints on effect sizes while allowing the data to refine the relationships between functional space and life stage classification. Finally, to regulate the complexity of the smooth terms in the model, we assigned Student's t priors to the standard deviations of the smoothing splines for both adults and juveniles. These priors allow for moderate variability while preventing extreme deviations, ensuring a balance between flexibility and regularization. Moreover, they provide robustness against potential outliers while still permitting the estimation of smooth, biologically meaningful relationships between the functional space dimensions and life stage classification.

To assess the impact of prior specification on model inference, we ran a second model using weakly informative priors automatically set based on the data and model structure. We compared the two models using leave-one-out cross-validation (LOO) and the expected log predictive density criterion (ELPD) to assess model fit and predictive performance. Differences in posterior distributions and parameter estimates were also examined to evaluate the influence of prior specification.

*Mapping life stages in the functional space*

To evaluate the relationship between the functional space of *Posidonia oceanica* fish assemblage and life stage classification, we used the fitted Bayesian model to predict probabilities across the functional space. Specifically, we applied the 'fitted' function from the *brms* package to a new data grid representing the functional space derived from the traits' probability density function (TPD, see main text). This approach allowed us to estimate the probability of species being classified as either adults, juveniles and both (adults + juveniles) based on their functional trait positions, providing a probability surface reflecting how the likelihood of each life stage varied across the functional space.

Below, a slightly modified version of part of the original *funspaceGAM* code adapted to a Bayesian framework. We also provided the results from *brms* (brm_model.rds) and *funspace* (funspace_data.rds) as a R Data Serialization file (.rds), to speed-up the code and avoiding excessive computation time. All supplementary files can be found here: https://figshare.com/s/385ded19fbcda07f9f5c

```
#**********************************************************************************************
*
# The following code is partially extracted from the original function
# funspaceGAM in the funspace package. The code has been rearranged to allow
# the use of a Bayesian multinomial regression with splines into the original
# function.

#*****************************************Load
models*******************************************
# Load the brms model results (to speed up)
brm_model<- readRDS("brm_model.rds")
# Load functional space analysis
funspace_data = readRDS("funspace_data.rds")

#**********************************Predict across the whole trait
space*********************************
predGrid <- funspace_data$parameters$evaluation_grid
colnames(predGrid) <- c("FAMD1", "FAMD2") # Set column names to match with the brm_model
predAux <- as.data.frame(fitted(object = brm_model,newdata = predGrid,scale = "response")) # Takes a lot
of time!
grid_prediction = function(prediction,se_prediction){
funspace_data$global$predicted <- cbind(funspace_data$parameters$evaluation_grid,
                          predicted = prediction,
                          se = se_prediction)
                          imagePred <- imageSEPred <- funspace_data$global$images$TPD.quantiles
                          imagePred[ , ] <- imageSEPred[ , ] <- NA
                          for(k in 1:length(unique(funspace_data$parameters$evaluation_grid[, 2]))){
                          colAux <- subset(funspace_data$global$predicted,
                          funspace_data$global$predicted[,2] ==
    unique(funspace_data$parameters$evaluation_grid[, 2])[k])
                          imagePred[, k] <- colAux$predicted
```

```
                                imageSEPred[, k] <- colAux$se
  }
  ### apply mask
  maskQuant <- funspace_data$global$images$TPD.quantiles
  maskQuant[!is.na(maskQuant)] <- 1
  imagePredMask <- imagePred * maskQuant
  imageSEPredMask <- imageSEPred * maskQuant
  funspace_data$global$images$predicted <- imagePredMask
  funspace_data$global$images$SE.predicted <- imageSEPredMask
  imagePredMask
}

#*************************Extract the conditional probability for each life
    stage***************************
Adults = grid_prediction(prediction = predAux$`Estimate.P(Y = Adult)`,
                    se_prediction = predAux$`Est.Error.P(Y = Adult)`)
Juveniles = grid_prediction(prediction = predAux$`Estimate.P(Y = Juvenile)`,
                      se_prediction = predAux$`Est.Error.P(Y = Juvenile)`)
Both = grid_prediction(prediction = predAux$`Estimate.P(Y = Both)`,
                  se_prediction = predAux$`Est.Error.P(Y = Both)`)

#**************************************Plot the
    result******************************************
par(pty="s",mfrow=c(1,3), las = 1)
pch_points = c(22,24,21)[as.numeric(as.factor(posidonia_fish$feeding_behaviour_original))]
color_gradient <- colorRampPalette(c("red","yellow","green"))(100)
image(x = as.numeric(rownames(Juveniles)),
    y = as.numeric(colnames(Juveniles)),
    z = Juveniles, col = color_gradient, zlim = c(0,1), cex.axis=1.5, xlab = "", ylab = "")
contour(x = as.numeric(rownames(Juveniles)),
      y = as.numeric(colnames(Juveniles)),
      z = Juveniles, add = TRUE, col = "black", lwd = .75)
points(posidonia_fish[,2:3], pch = pch_points, col = "black",bg = "lightgray")
abline(h = 0, v = 0, lty = 2)

image(x = as.numeric(rownames(Both)),
    y = as.numeric(colnames(Both)),
    z = Both, col = color_gradient, cex.axis=1.5, xlab = "", ylab = "", zlim = c(0,1))
contour(x = as.numeric(rownames(Both)),
      y = as.numeric(colnames(Both)),
      z = Both, add = TRUE, col = "black", lwd = .75)
points(posidonia_fish[,2:3], pch = pch_points, col = "black",bg = "lightgray")
abline(h = 0, v = 0, lty = 2)

image(x = as.numeric(rownames(Adults)),
    y = as.numeric(colnames(Adults)),
    z = Adults, col = color_gradient, cex.axis=1.5, xlab = "", ylab = "", zlim = c(0,1))
contour(x = as.numeric(rownames(Adults)),
      y = as.numeric(colnames(Adults)),
      z = Adults, add = TRUE, col = "black", lwd = .75)
points(posidonia_fish[,2:3], pch = pch_points, col = "black",bg = "lightgray")
abline(h = 0, v = 0, lty = 2)
```

#**************************************************END*******************************************
**

*Bayesian generalized additive model results*

The difference in predictive accuracy, as indexed by the expected log predictive density (ELPD-LOO), indicated that the model with informative priors had a superior predictive accuracy (ELPD = -166.80; diff-ELPD = -0.90 +- 1.98, p = 0.648), suggesting that incorporating prior knowledge improved model performance compared to relying solely on default (weakly informative) priors.

The model revealed significant associations between the predictors (FAMD1 and FAMD2) and the likelihood of individuals being classified as either adults or juveniles (Table S1). The estimated smooth term standard deviations (sds) suggest notable variation in the effects of these predictors across categories. For the adult category, the standard deviation of the smooth effect of FAMD1 was 1.12 (95% CI: 0.16 - 2.10), while for FAMD2, it was 1.13 (95% CI: 0.13 - 2.11). In contrast, for the juvenile category, the estimated smooth effects were -1.03 (95% CI: -1.99 - -0.07) for FAMD1 and -0.94 (95% CI: -1.91 - 0.01) for FAMD2, indicating a negative association with juvenile classification.

The regression coefficients further support these trends. The intercept for adults was -0.44 (95% CI: -0.78 - -0.09) and -0.71 (95% CI: -1.06 - -0.35) for juveniles, suggesting that the reference category both (adults + juveniles), was the most frequent. The effect of FAMD1 on adults was positive (1.12, 95% CI: 0.16 - 2.10), whereas it was negative for juveniles (-1.03, 95% CI: -1.99 - -0.07). Similarly, FAMD2 had a positive effect on adults (1.13, 95% CI: 0.13 - 2.11) but a negative effect on juveniles (-0.94, 95% CI: -1.91 - 0.01), though the confidence interval for the latter slightly overlapped zero.

All parameters showed strong convergence diagnostics ($\widehat{R} = 1$), with high effective sample sizes ($ESS_{Bulk} > 5,400$, $ESS_{Tail} > 2,400$), indicating robust and reliable posterior estimates. Overall, these results suggest that FAMD1 and FAMD2 play differential roles in the life stage classification, with stronger positive effects for adults and negative effects for juveniles.

**Table S1** - Summary of parameter estimates for the Bayesian generalized additive model with smoothing splines. The table includes the posterior mean estimate (Estimates), standard error (SE), and the 95% credible interval (l-95% CI, u-95% CI) for each parameter. $\hat{R}$ is the potential scale reduction factor (aka Gelman-Rubin diagnostic), with values close to 1 indicating optimal convergence. $ESS_{Bulk}$ and $ESS_{Tail}$ represent the effective sample sizes for bulk and tail posterior distributions, respectively. $\mu_{Adult}$ and $\mu_{Juvenile}$ correspond to the log-odds of the corresponding categories relative to the reference one (both, adults + juveniles). The predictors s(FAMD1) and s(FAMD2) are smooth functions of the latent dimensions from the factor analysis of mixed data (FAMD), while sds indicates the standard deviation of the smoothing spline coefficients for each predictor and category. These values indicate the amount of variation in the smooth function's contribution to the model.

| Smoothing Spline Hyperparameters | | | | | | | |
|---|---|---|---|---|---|---|---|
| | Estimates | SE | l-95% CI | u-95% CI | $\hat{R}$ | $ESS_{Bulk}$ | $ESS_{Tail}$ |
| sds[$\mu_{Adult}$_s(FAMD1)] | 1.12 | 0.5 | 0.16 | 2.10 | 1 | 5,541 | 3,413 |
| sds[$\mu_{Adult}$_s(FAMD2)] | 1.13 | 0.51 | 0.13 | 2.11 | 1 | 5,530 | 3,121 |
| sds[$\mu_{Juvenile}$_s(FAMD1)] | -1.03 | 0.49 | -1.99 | -0.07 | 1 | 6,735 | 2,822 |
| sds[$\mu_{Juvenile}$_s(FAMD2)] | -0.94 | 0.5 | -1.91 | 0.01 | 1 | 5,412 | 2,443 |

| Regression coefficients | | | | | | | |
|---|---|---|---|---|---|---|---|
| | Estimates | SE | l-95% CI | u-95% CI | $\hat{R}$ | $ESS_{Bulk}$ | $ESS_{Tail}$ |
| $\mu_{Adult}$_Intercept | -0.44 | 0.18 | -0.78 | -0.09 | 1 | 5,459 | 3,222 |
| $\mu_{Juvenile}$_Intercept | -0.71 | 0.19 | -1.06 | -0.35 | 1 | 5,351 | 2,759 |
| $\mu_{Adult}$_s(FAMD1) | 1.12 | 0.5 | 0.16 | 2.10 | 1 | 5,541 | 3,413 |
| $\mu_{Adult}$_s(FAMD2) | 1.13 | 0.51 | 0.13 | 2.11 | 1 | 5,530 | 3,121 |
| $\mu_{Juvenile}$_s(FAMD1) | -1.03 | 0.49 | -1.99 | -0.07 | 1 | 6,735 | 2,822 |
| $\mu_{Juvenile}$_s(FAMD2) | -0.94 | 0.5 | -1.91 | 0.01 | 1 | 5,412 | 2,443 |

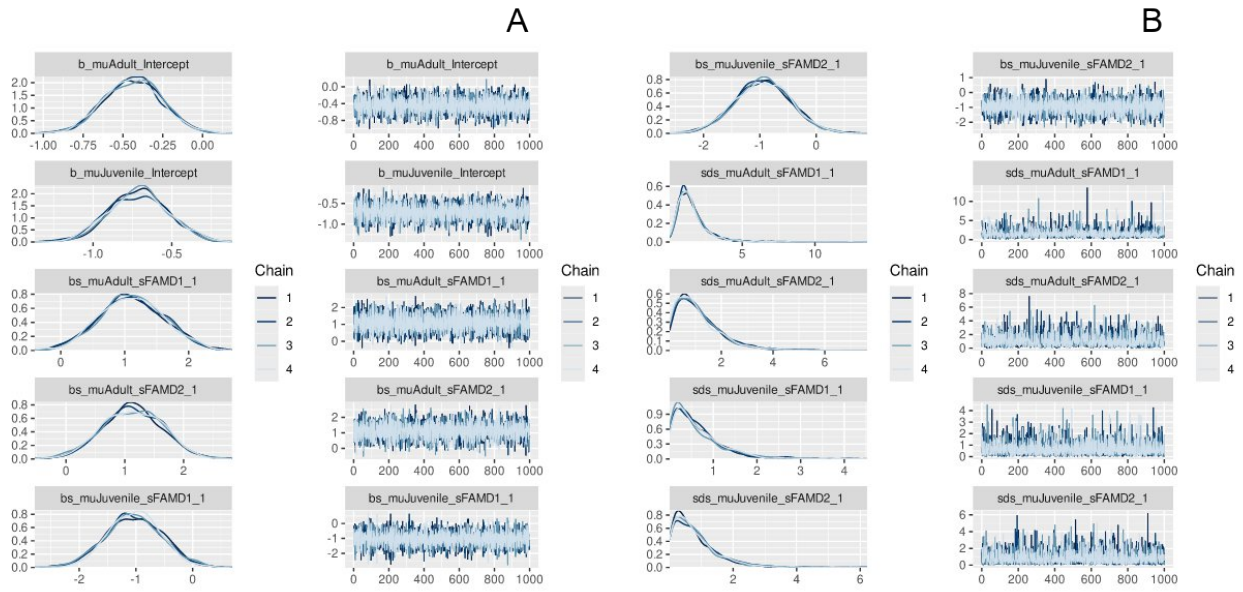


**Figure S2** - Posterior distributions and trace plots for the parameters of the Bayesian multinomial regression with smoothing splines. Panel A displays the posterior distributions (left) and trace plots (right) for the intercepts and smooth terms. Panel B shows the corresponding posterior distributions and trace plots for the smoothing standard deviations (sds) associated with each smooth term. The consistent mixing of chains in the trace plots and the smooth posterior distributions indicate convergence and effective sampling across all parameters.

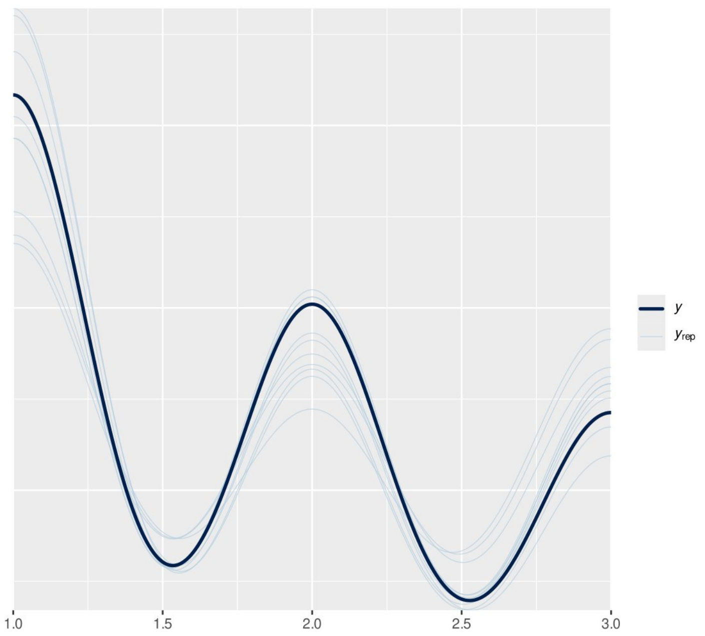


**Figure S3** - Posterior predictive check for the Bayesian generalized additive model. The observed data (y, dark blue line) is overlaid with replicated data ($y_{rep}$, light blue lines) generated from the posterior predictive distribution. The close alignment between the observed and replicated data indicates that the model captures the key patterns in the data and provides a good fit.

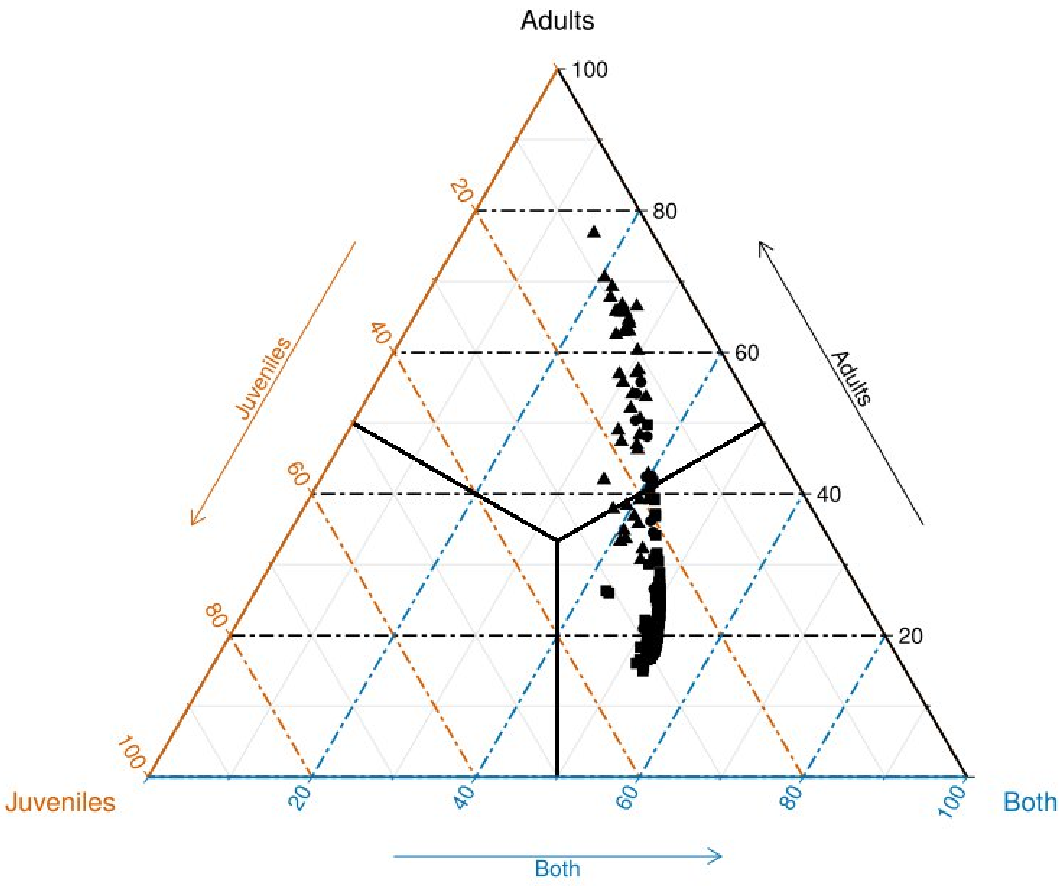


**Figure S4** - Ternary plot illustrating the proportional distribution of fish across life stage categories: juveniles, adults, and both (species occurring in both stages). Point coordinates in the plot correspond to the predicted probability of the Bayesian multinomial regression with smoothing splines. Triangles, circles and squares are for active predators, grazers and ambush predators, respectively. The black lines represent classification thresholds (50%) separating different life stage composition zones based on the proportions of juveniles, adults, and species occurring in both stages. Thresholds likely indicate specific regions where the dominance of one life stage over the others shifts. Grid lines indicate percentage values along each axis, with orange lines representing increasing juvenile proportions, blue increasing presence in both stages, and black increasing adult proportions. The plot highlights a predominant clustering of observations toward the both-dominated region, suggesting a non-random distribution of life stages. Almost all active predators fall within the adult-dominated region, and no species within the juvenile-dominated region.
